# Biosynthesis of Arcyriaflavin F from *Streptomyces venezuelae* ATCC 10712

**DOI:** 10.1101/2024.04.17.589956

**Authors:** Hung-En Lai, Lewis Tanner, Agata Kennedy, Soo Mei Chee, Paul S Freemont, Simon J Moore

## Abstract

Indolocarbazoles are natural products with a broad spectrum of reported bioactivities. A distinct feature of indolocarbazole biosynthesis is the modification of the indole and maleimide rings by regioselective tailoring enzymes. Here, we study a new indolocarbazole variant, which is encoded by the *acfXODCP* genes from *Streptomyces venezuelae* ATCC 10712. First, we characterise this pathway by expressing the *acfXODCP* genes in *Streptomyces coelicolor*, which led to the production of a C-5/C-5’-dihydroxylated indolocarbazole. We name this new product arcyriaflavin F. Second, we demonstrate the flavin-dependent monooxygenase AcfX catalyses the C-5/C-5’ dihydroxylation of the unsubstituted arcyriaflavin A into arcyriaflavin F. Interestingly, AcfX shares homology to EspX from erdasporine A biosynthesis, which instead catalyses a single C-6 indolocarbazole hydroxylation. In summary, we report a new indolocarbazole biosynthetic pathway and a regioselective C-5 indole ring tailoring enzyme AcfX.

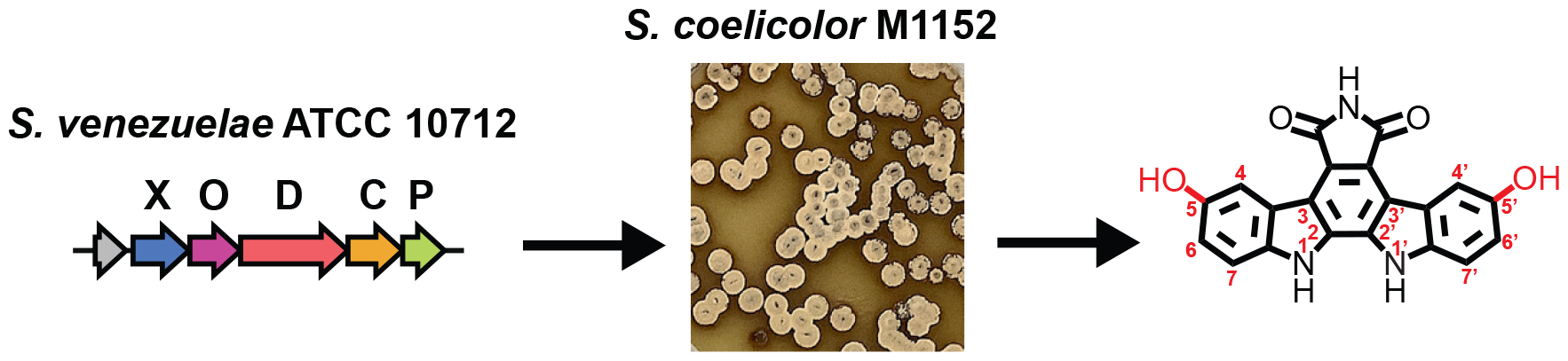

## Introduction

Bisindoles and indolocarbazoles are a diverse natural product family with potential therapeutic applications, including Gram-positive antibacterials^[1–3]^, antitumor^[4–6]^, antifungal and anti-malaria bioactivity^[7]^. Midostaurin (Rydapt®, Novartis) is an approved glycosylated indolocarbazole for clinical treatment of acute myeloid leukemia^[8]^. Two natural product indolocarbazoles, staurosporine and rebeccamycin, specifically inhibit protein kinases^[9]^ and DNA topoisomerase I^[5]^, respectively. As an indolocarbazole sub-class, the arcyriaflavins were originally isolated from the slime mold species *Arcyria denudate* and thus a prominent pigment within the red fruiting bodies of this organism. Arcyriaflavin A (**1**) has a core indolocarbazole scaffold fused to a maleimide ring. Although evolutionarily distinct, these natural products are also made by bacteria. Bacterial biosynthesis of indolocarbazoles is either low or variable, therefore either synthetic chemistry^[10–14]^ or recombinant approaches are used to make these molecules. A number of indolocarbazole variants were recombinantly biosynthesised from metagenomic sources by the Brady group^[15–18]^, while indolocarbazoles have been overproduced in *Pseudomonas putida*^[19]^ and *Streptomyces albus* J1074^[20]^.

The biosynthesis of indolocarbazoles was first characterised for the rebeccamycin pathway^[6,21,22]^. The indolocarbazole scaffold forming enzymes are RebO, -D, -C and -P, with the last letter of the protein name shared in all indolocarbazole pathways – i.e., for staurosporine biosynthesis: StaO, -D, -C and -P (**Figure 1A**). Indolocarbazole biosynthesis requires two L-tryptophan substrates. The flavin adenine dinucleotide (FAD)-dependent L-tryptophan (L-Trp) oxidase (RebO) forms an indole-3-pyruvate imine, which binds to a haem-containing oxygenase (RebD) and reacts with an enamine L-Trp intermediate^[23,24]^. A highly valent Fe(III)-O centre within the haem group catalyses single electron abstraction from the β carbon position to create a radical, which can close to form a new C-C bond, followed by a second H abstraction with Fe(IV)-OH and double bond rearrangement to provide a new pyrrole ring and the product chromopyrrolic acid (CPA)^[21,23]^. Last, a P450 enzyme RebP, catalyses a similar radical mechanism to form a new C-C bond^[25]^. RebP acts in unison with a FAD-dependent enzyme (RebC) to catalyse an 8-electron oxidation of two carboxylic acids to a maleimide ring^[22]^, releasing CO_2_, H_2_O and NADH as products, creating arcyriaflavin A (**Figure 1B**). Variant indolocarbazoles stem from modifications to the indole ring or the maleimide ring^[15,26,27]^. Examples include 2-pyrrolidinone indolocarbazoles like K-252c and carboxypyrrole indolocarbazole erdasporine A, which were isolated from soil-derived sources^[15]^. Other hydroxylated variants include BE-13793C^[5]^, arcyriaflavin E^[28]^ and hydroxysporine^[16]^. These findings emphasize the role of late stage regioselective tailoring enzymes that decorate the indole ring. This includes the flavin-dependent C-6 hydroxylase EspX^[15]^ and a two-component cytochrome P450 C-5 hydroxylase HysX1/X2^[16]^. There are also a range of indolocarbazole variants isolated from environmental Actinobacteria^[28–31]^.

**Figure 1.**
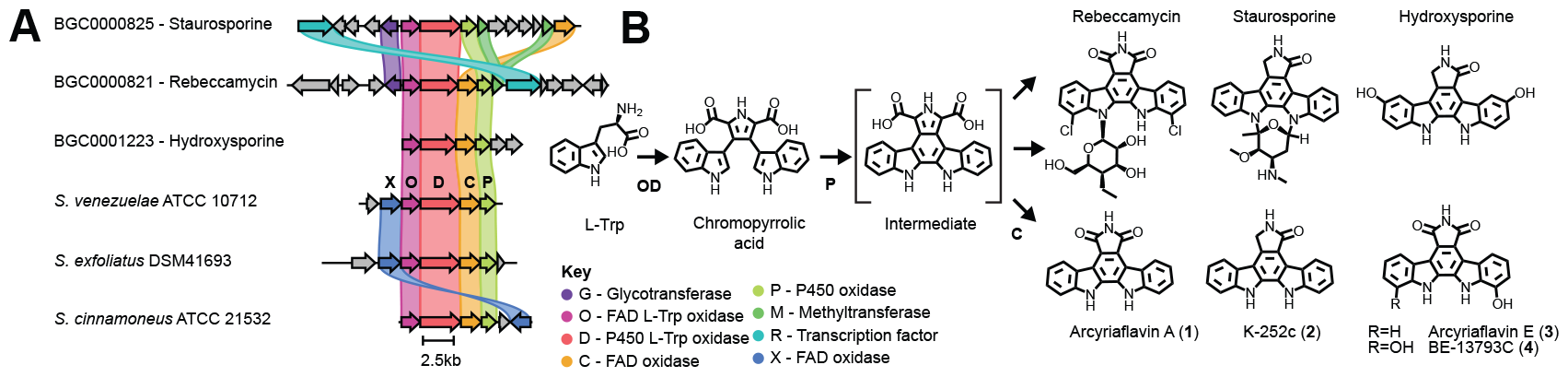
Overview of indolocarbazole biosynthesis and genetic organisation of the *S. venezuelae* ATCC 10712 arcyriaflavin F BGC. **A**: Alignment of selected known (with MIBiG IDs) and putative indolocarbazole biosynthetic gene clusters using Clinker software^[34]^. **B**: A general scheme for indolocarbazole biosynthesis and a selection of natural products. As an exception the rebeccamycin biosynthesis begins with C-7 chlorination of L-Trp, whereas other indole tailoring steps occur late in the biosynthetic pathway.

Building upon our recent interest in *Streptomyces venezuelae* ATCC 10712^[32,33]^, we identified a five-gene indolocarbazole biosynthetic gene cluster (BGC) on its chromosome, which shares similarity to rebeccamycin (MIBiG: BGC0000821) and staurosporine (MIBiG: BGC0000825) BGCs^[6,21]^ (**Figure 1A**). Distinctly, the *S. venezuelae* indolocarbazole BGC carries an uncharacterised flavoprotein (Locus tag: VNZ_RS03650) with similarity to EspX from erdasporine A biosynthesis. We report the characterisation of the *S. venezuelae* indolocarbazole BGC, the flavoprotein enzyme and the pathway’s product arcyriaflavin F.

## Results and Discussion

### Characterisation of the *S. venezuelae* arcyriaflavin BGC

*S. venezuelae* ATCC 10712 encodes 30 biosynthetic gene clusters (BGCs) as predicted by AntiSMASH bioinformatics analysis^[35]^. Five of these BGCs are known to encode chloramphenicol^[36,37]^, jadomycin^[36]^, watasemycin^[38]^, venepeptide^[39]^ and pikromycin^[40]^ biosynthesis. The other 25 BGCs remain uncharacterised, with one encoding five genes homologous to *rebO, -D, C* and *-P* from rebeccamycin biosynthesis, as well as *espX* from the erdasporine A pathway (MIBiG: BGC0001336). Interestingly, *S. venezuelae* was previously reported to make a form of arcyriaflavin^[41]^. However, production was only detected after extensive genetic manipulation of the strain, the details of which were not subsequently published (Professor Mervyn Bibb, personal communication). To investigate whether a laboratory condition could trigger the *S. venezuelae* ATCC 10712 type strain to make arcyriaflavin A or variants, we initially screened a range of routine *Streptomyces* growth media^[42]^, using both liquid and solid agar cultures. Analysis of organic extracts separated by C18 reverse-phase high-performance liquid chromatography (HPLC) failed to detect any arcyriaflavin-like metabolites by either ultraviolet-visible (UV-Vis) absorbance or mass spectrometry (MS) detection approaches, when compared to an arcyriaflavin A standard. This suggested that cluster activation requires either an unknown effector, likely present in the environment, or through genetic modification.

To investigate the proposed *S. venezuelae* arcyriaflavin (*acfXODCP*) BGC, we switched to heterologous gene expression. Using Gibson DNA assembly, we inserted the *acfXODCP* or *acfODCP* genes within the integrative pAV-gapdh vector^[43,44]^, which carries a strong P_*gapdh(EL)*_ constitutive promoter upstream of the operon and a *phiC31* integrase for integration into the *S. coelicolor* M1152 chromosome^[44]^. An empty vector (pAV-gapdh) lacking the *acf* BGC was also included as a control. After conjugation with *S. coelicolor* M1152, exconjugants from the recombinant *acfXODCP* or *acfODCP* strains showed production of a light brown or orange pigment, respectively, after approximately 7-10 days of growth. The control colonies were a typical white-grey colouration, suggesting that the recombinant arcyriaflavin BGCs were active and responsible for pigment production. Next, we analysed ethyl acetate extracts of solid MS agar growth of the *S. coelicolor* M1152 recombinant *acfODCP* and *acfXODCP* strains by C-18 high-performance liquid chromatography (HPLC) coupled with mass spectrometry (MS) in positive ion mode. First, no apparent indolocarbazoles or intermediates were detected from extracts of the M1152 control strain. For the *acfODCP* strain, known indolocarbazole biosynthetic intermediates and side-products were detected in low quantities, with the exception that chromopyrrolic acid (CPA) (Expected *m/z* = 384.10) was absent. CPA was previously detected as an intermediate during heterologous production of rebeccamycin in *Streptomyces albus*^[45]^. An arcyriaflavin A standard had a retention time (RT) of 39.9 min with an [M+H]^+^ *m/z* = 326.09 **(1)**. In the *acfODCP* strain, detected indolocarbazoles included arcyriaflavin A (**1**) at [M+H]^+^ *m/z* = 326.09 (RT = 39.9 min), K-252c (also known as staurosporinone) (**2**) at [M+H]^+^ *m/z* = 312.11 (RT = 35.5 min) and 7-hydroxy-K-252c (or hydroxystaurosporinone) (**3**) at [M+H]^+^ *m/z* = 328.11 (RT = 33.75 min). In the *acfODCP* pathway variant, the major product was arcyriaflavin A (**1**), constituting over 95% of the total extracted ion chromatogram (EIC) for all indolocarbazole metabolic species detected (**Figure 2**). The putative species corresponding to K-252c (**2**) and 7-hydroxy-K-252c (**3**) accounted for <5% of the total EIC. For the *acfXODCP* strain, arcyriaflavin A was absent. Instead, new peaks for a minor (∼2%) and a major (∼98%) product with [M+H]^+^ *m/z* = 342.08 (RT = 32.8 min) and 358.08 (RT = 24.7 min) was observed, respectively. These products corresponded to the addition of 15.99 and 31.99 mass units, or one and two oxygen atoms respectively, suggesting potential hydroxylation. The decrease in RT for both these molecules in comparison to arcyriaflavin A (compound **1**), also suggested reduced hydrophobicity. For this, we initially assigned these peaks as potential mono- and dihydroxylated analogues of arcyriaflavin A (**1a** and **1b**), respectively. Interestingly, the dihydroxylated species at 24.7 min, consistently split into three peaks both on Ultraviolet-visible light (UV-Vis) absorbance and MS (EIC of 358.08) detection. Previous chemical synthesis of arcyriaflavin analogues, including arcyriaflavin F, was reported, which noted general instability for the hydroxylated species^[4]^. Overall, the HPLC-MS data supports that arcyriaflavin A is the major product for the recombinant strains expressing the *acfODCP* genes, whereas dihydroxylated analogue of arcyriaflavin A, is the major product for the *acfXODCP* pathway.

**Figure 2.**
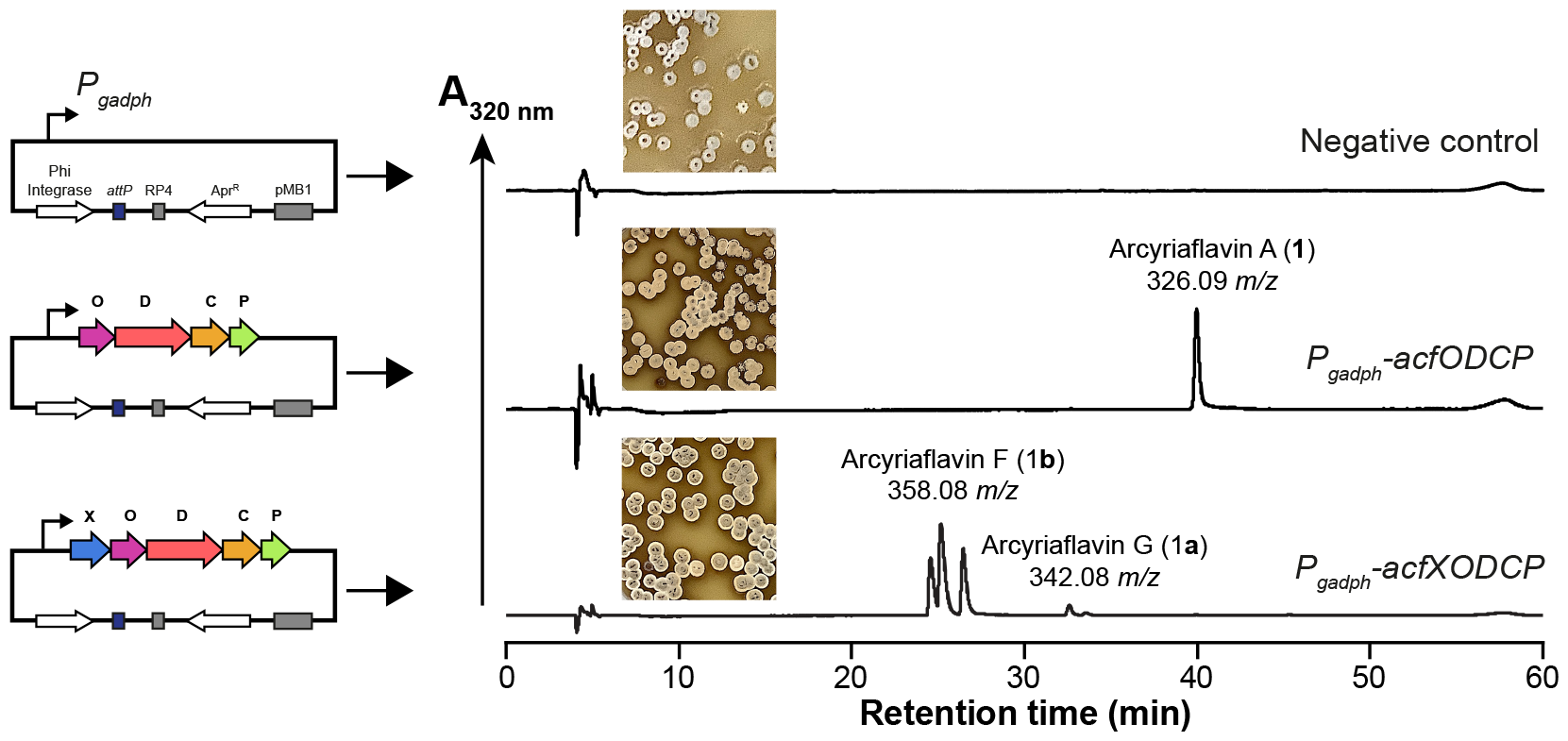
HPLC chromatograms of organic extracts of *S. coelicolor* M1152 overexpressing the *S. venezuelae* ATCC 10712 *acfODCP* and *acfXODCP* operons. Cells were grown for 7 days at 30°C on MS agar plates and prepared for HPLC-MS analysis as described in the methods. Chromatograms represent relative visible light absorbance at 320 nm, with the dominant mass for each peak displayed. Extracted ion chromatograms are provided in Figure S1. One representative HPLC chromatogram is shown from three independent experiments. An analytical standard of **1** (Santa-Cruz chemicals) was run in parallel to confirm the *acfODCP* product.

### Structural characterisation of arcyriaflavin F and C-5/C-5’ hydroxylation

The product of the *acfXODCP* recombinant pathway is consistent with mono- (minor) and dihydroxylated (major) variants of arcyriaflavin. To confirm the structure, we scaled-up the growth of the *S. coelicolor* M1152 *acfXODCP* strain (in 4L of MR5 media) for compound isolation and NMR analysis. Due to low quantities of the presumed mono-hydroxylated analogue arcyriaflavin G (**1a**), we did not isolate this species for structural characterisation. Next, we purified the major product of the *S. coelicolor* M1152 *acfXODCP* strain as outlined in the methods. From 4 litres of culture ferment, 13.8 mg of **1b** was purified as a brown solid. Based on the EIC for the mass 358.08 *m/z* and related pathway metabolites, the purity of **1b** was estimated to be greater than 99%. **1b** was dissolved in 600 μL d_6_-DMSO and the ^1^H, ^13^C, HMBC, HMQC, HSQC and ROESY NMR spectra were obtained (NMR data is provided in **Table S2** and **Figure S2**). The ^1^H NMR spectrum showed 11 protons, including three aromatic protons between δ 7.02 (dd), 7.59 (d) to 8.40 (d). In comparison to previous NMR spectra for arcyriaflavin A, four aromatic protons (between δ 7.35 and 8.98) were previously observed for the C-4 and C-7 positions^[28]^, suggesting one of the positions is connected to a new group. Further, there were three downfield-shifted protons, which were assigned as a single OH (δ 9.2) and two NH (δ 10.85 and 11.4) groups. For the ^13^C NMR spectrum, 20 carbons were present, consistent with the arcyriaflavin A structure^[28]^. Based on the combined MS and NMR spectra, the data supports the structure of **1b** with two hydroxyl groups at the C-5 and C-5’ indole ring positions (atomic position 10/22 in **Figure 2** and **Table S2**). The C-5/C-5’ hydroxyl group assignment is supported by the following observations: ROESY through-space interactions between the indole NH group and the C-7 proton, and ROESY connections for the C-5/C-5’ hydroxyl proton with the C-4 and C-6 protons. In addition, C-5 is downfield shifted (151.48 ppm) in comparison to C-4 (108.82 ppm) and C-6 (116.32 ppm), which favours C-5/C-5’ as the hydroxylation position. In addition, the HSQC and HMQC spectra for arcyriaflavin F also revealed three sets of C-H couplings each, indicating mixed species. This is consistent with the split HPLC-MS peak and previous literature for chemical synthesis of arcyriaflavin F^[4]^. Based on the major species, we assign **1b** as a C-5/C-5’ dihydroxylated form of arcyriaflavin,

### AcfX catalyses C-5 hydroxylation of arcyriaflavin A

C-5 and C-5’ indole hydroxylation is observed in hydroxysporine biosynthesis, but in contrast, this reaction is distinctly catalysed by two HysX1 and HysX2 cytochrome P_450_ enzymes^[16]^. AcfX shares 54.7% identity to EspX, a flavin adenine dinucleotide (FAD) hydroxylase that catalyses a single hydroxylation at the C-6 indole ring position to form erdasporine A biosynthesis^[15]^. To study AcfX enzymatic activity, we cloned *acfX* with an C-terminal His_6_-tag into pSF1C-SP44 plasmid. Then we used *E. coli* Rosetta (DE3) pLysS-Rare as recombinant host to overproduce AcfX for immobilized metal ion affinity (IMAC) purification (see methods). AcfX-His_6_ was observed as a single band at approximately 67 kDa by denaturing polyacrylamide gel electrophoresis. During the purification, AcfX purified as pale-yellow coloured protein and was generally unstable in a range of common buffer/salt conditions. The best condition we could establish was the S30 buffer (see methods), with the addition of 15% (v/v) glycerol to stabilise the protein. However, AcfX also gradually precipitated over prolonged incubation and was generally found to be unstable. In terms of the yellow pigment, and prediction that AcfX is a flavoprotein, we measured the UV-Vis light absorption spectrum and observed major absorbance peaks at 350 nm and 420 nm (**Figure 4B**), typical of flavin bound to the protein. This was further confirmed using FAD and FMN standards by HPLC-MS analysis (**Figure 4C**).

**Figure 3.**
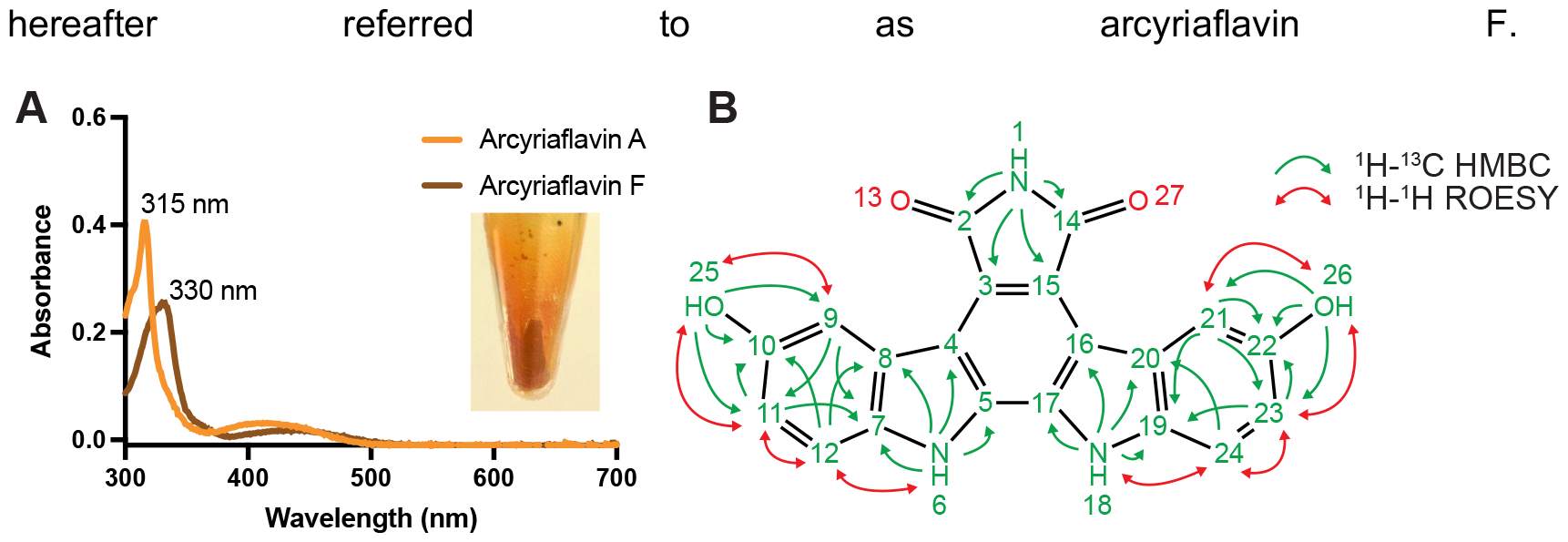
Characterisation of arcyriaflavin A. **A**: UV-Viv absorbance spectrum of purified arcyriaflavin A and arcyriaflavin F in ethanol (inset image – dried arcyriaflavin F extract). **B**: Structure of arcyriaflavin F and ^1^H-^13^C HMBC and ^1^H-^1^H ROESY correlations. 1D and 2D NMR spectra are provided in Figure S2. Atom positions are labelled according to Table S2.

**Figure 4.**
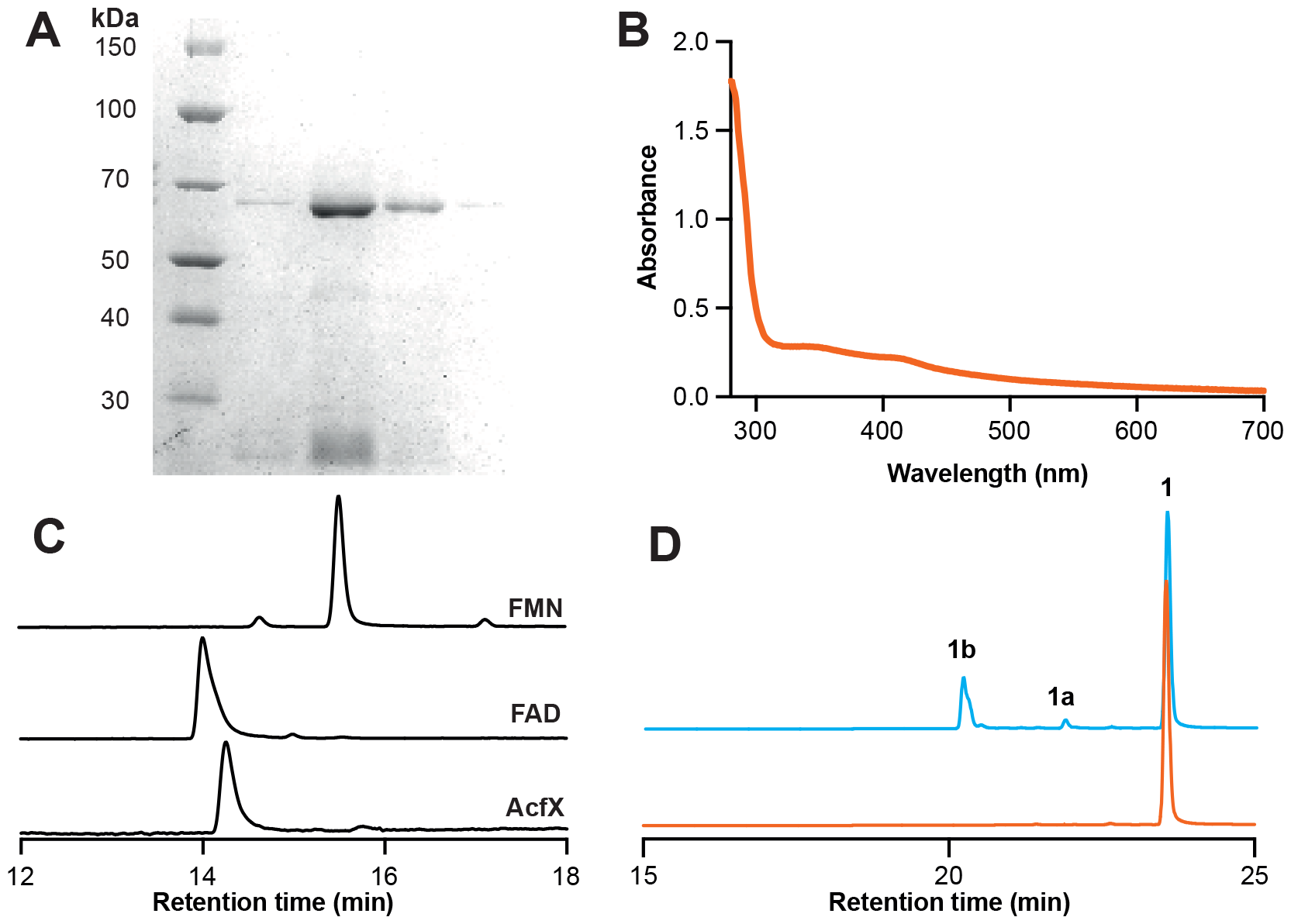
Characterisation of the AcfX C-5/C-5’ indolocarbazole hydroxylase. **A**: Denaturing polyacrylamide gel electrophoresis (PAGE) of purified AcfX. **B**: UV-Visible absorbance of AcfX. **C**: HPLC analysis of AcfX flavin cofactor with analytical standards of FMN and FAD. **D**: HPLC-MS analysis of an AcfX reaction after 16 hours of incubation. Negative control and time course incubations are shown in Figure S3. The assay and controls were repeated on four independent experiments.

To study AcfX enzymatic activity, we used C-18 HPLC analysis (Method B) to monitor conversion of **1** into **1b**. 10 μM of purified AcfX was incubated in the presence of 50 μM **1**. To provide an artificial source of electrons, we supplemented the reaction with 1 mM NADPH. After 16 hours of incubation, approximately 25% of **1** (RT = 20.7 min) was converted into **1b** (RT = 23.5 min) (**Figure 4D**). Interestingly, trace levels (3%) of the mono-hydroxylated **1a** were also detected (RT = 21.87 min), which complements the cell-based production data (**Figure 2**). For the controls, in the absence of AcfX or NADPH, no product conversion was observed (**Figure S3**). This reaction was repeated on four independent experiments, where the yield varied between 5-30% conversion (**Figure S3**). While the enzymatic activity of AcfX was modest, our data supports the role of AcfX as a FAD-dependent C-5/C-5’ arcyriaflavin A hydroxylase. We suggest that NADPH is sufficient to reduce the AcfX-bound FAD cofactor to FADH, prior to oxygen splitting and FAD-4a-OOH formation, whereas the reaction is likely more favourable and proceeds to completion (as observed in Figure 2) in cells via a non-specific reductase partner. FAD monooxygenases are known to readily hydroxylate kinetically accessible carbanions, which are found widely in indole, phenols, and pyrrole rings. One example is OxyS from oxytetracycline biosynthesis, which shares 33% identity to AcfX. OxyS modifies the enol and keto scaffold in the final steps of oxytetracycline biosynthesis^[46]^.

### Conclusions

Herein, we study the arcyriaflavin F BGC from *S. venezuelae*. We show MS and NMR data supporting C-5/C-5’ hydroxylation at the indole ring positions. This tailoring reaction is distinct to the related erdasporine A, which has only a single hydroxyl group at the C-6 position catalysed by the EspX hydroxylase^[15]^. The biosynthetic rationale of AcfX is also distinct to hydroxysporine biosynthesis, which installs C-5/C-5’ hydroxyl groups through the action of a dual cytochrome P_450_ (HysX1 and HysX2) system. Our work reveals a new insight into regioselective tailoring of bisindoles and indolocarbazole natural product biosynthetic pathways and contributes to the growing area of synthetic biology for natural products.

## Experimental Section

### Bacterial strains and growth media

For general cloning and plasmid purification, *E. coli* JM109 or DH10β strains were used. *E. coli* strains were grown in Luria-Bertani (LB) autoclaved media with appropriate antibiotic concentrations (apramycin 25 μg/mL, nalidixic acid 30 μg/mL) at 30 °C or 37 °C. *S. venezuelae* ATCC 10712 cells (obtained from DSMZ Cat. DSM-40230) were grown in GYM/LB/MS/Potato agar media at 30 °C. *S. coelicolor* M1152 (provided by Prof. Mervyn Bibb, John Innes Centre, UK) was grown in modified R5 (MR5) media^[47]^ (0.25 g/L K_2_SO_4_, 10.12 g/L MgCl_2_.6H_2_O, 10.0 g/L D-glucose, 0.1 g/L casamino acids, 5.0 g/L yeast extract, 2.0 g/L CaCO_3_, 0.2% (w/v) trace elements, adjusted to pH 6.8, and grown at 30 °C with shaking (150-200 rpm).

### Assembly and conjugation of the *acf* BGC

The *acf* BGC was amplified from *S. venezuelae* ATCC 10712 genomic DNA as ∼3 kb fragments and assembled into the pAV-gapdh vector^[43,44]^ (a gift from Professor Jay Keasling) by Gibson DNA assembly. Each PCR fragment was generated as a PCR product using Q5® High-Fidelity DNA Polymerase and ensuring ∼30 bp overlap between the homologous regions. 5 μL of NEBuilder® HiFi DNA Assembly Master Mix (2X) was added to 5 μL of DNA mixture containing 0.05 pmol of each PCR fragment and incubated at 50 °C for 2 h before transforming into competent *E. coli* DH10β cells. The *acf* BGC was then verified by commercial long-read sequencing (by Full Circle Labs, UK). Oligonucleotides and plasmids are listed in Table S1. Conjugation was performed using *E. coli* ET12567::pUZ002 into *S. coelicolor* M1152 as described previously^[42]^.

### HPLC-MS analysis

A single *S. coelicolor* M1152 colony was picked and inoculated into 10 mL autoclaved MR5 media with nalidixic acid and apramycin in 50 mL sterile tubes. Culture was incubated at 30 °C for 12 days and extracted with 1 volume of ethyl acetate. The mixture was further incubated at 30 °C, 200 rpm shaking for 1 h, and the organic layer was transferred to a fresh tube, dried *in vacuo* and reconstituted in 300 μL of MeOH. Pathway metabolites were analysed by HPLC-MS (Method A) using an Agilent 1100 LC system connected to a Bruker micrOTOF II MS and detected with positive electrospray ionization (ESI), using an ACE 5 AQ C-18 column (2.1 x 250 mm, 5 μm particle size). For HPLC analysis, buffer A was water with 0.1% (v/v) trifluoroacetic acid and buffer B was acetonitrile with 0.1% (v/v) trifluoroacetic acid, for a total running time of 60 min including 10 min post-run time. Metabolites were detected at 320 nm, close to the maximal extinction coefficient for arcyriaflavins. 1-10 μL samples were injected, with a flow rate of 0.2 mL/min and the following gradient: 0 min 5% B, 50 min 100% B, 55 min 100% B, 60 min 5% B (Method A). Absorbance spectra was monitored by Diode Array UV-Vis detection (Agilent Technologies).

### Structural characterisation of arcyriaflavin F

*S. coelicolor* M1152 exconjugant with the *acf* BGC integrated was inoculated into 4 L of MR5 media and grown for 12 d at 30 °C and 200 rpm. Culture was extracted with an equal volume of ethyl acetate, and the organic layer was separated and dried *in vacuo*. The crude extract was reconstituted in 4 mL of methanol and separated on a pre-equilibrated Biotage SNAP Ultra C-18 column 12 g attached to a Biotage Isolera Spektra system. Buffer A was ultrapure water (HPLC grade) and buffer B was acetonitrile (HPLC grade), for a total running time of 16.5 min including 2.4 min equilibration time. The column was run at 36 mL/min with the following gradient: 0 min 10% B, 2.35 min 15% B, sample loading, 6.60 min 45% B, 7.10 min 95% B with A as water and B as acetonitrile. Fractions that absorbed strongly at 290 nm and 335 nm were collected and pooled, dried *in vacuo* and the remaining solid was weighed. Approximately 7.8 mg of solid was dissolved in 600 μL of DMSO*-d*_6_ and transferred to an NMR tube 507 grade (Norell). The NMR sample was run on a Bruker Avance 400 MHz (DRX400) NMR spectrometer and data was collected for ^1^H, ^13^C, HMBC, HMQC, and HSQC experiments. NMR data and peak assignment were carried out using the MestReNova software 12.0.4-22023 release.

### Purification of AcfX and activity assay

*acfX*-His_6_ was PCR amplified by Q5 polymerase from *S. venezuelae* ATCC 10712 genomic data and cloned between NdeI and PacI in pSF1C-A-SP44^[32]^. Recombinant AcfX was over-produced in *E. coli* BL21 Gold (DE3) as a C-terminal His_6_-tagged protein. Cells were grown at 28°C, 200 rpm for 16 hours in autoinduction medium with 100 μg mL^-1^ carbenicillin. The day after, cells were collected by centrifugation at 6,000 × *g*, 4 °C for 20 min, and re-suspended in binding buffer (20 mM Tris-HCl pH 8, 500 mM NaCl, 5 mM imidazole) and lysed by sonication. Cell-lysates were clarified with centrifugation at 45,000 × *g*, 4 °C for 20 min and purified by gravity flow using nickel affinity chromatography (Cytiva). His_6_-tagged proteins were washed with increasing concentrations of imidazole (5, 30 and 70 mM) in 20 mM Tris-HCl pH 8, 500 mM NaCl and 10% glycerol, before elution at 400 mM imidazole. The purified protein was analysed by denaturing PAGE as described previously^[48]^. Purified proteins were buffer-exchanged and concentrated using a centrifugation filter (Amicon MWCO 10,000 Da) to a concentration of 2 mg/mL in 50 mM Tris-HCl pH 7.7, 14 mM Mg-glutamate, 60 mM K-glutamate and 15% (v/v) glycerol. Enzyme incubations were set up in buffer lacking glycerol, with different concentrations of arcyriaflavin A and NADPH, as specified in the results. Incubations were performed overnight at 30°C in a glass vial with 5% ethanol (v/v). Reactions were terminated with 1% (w/v) acetic acid, and centrifuged at 18,000 × *g* for 10 min, to remove precipitate. For HPLC analysis, buffer A was water with 0.1% (v/v) trifluoroacetic acid and buffer B was acetonitrile with 0.1% (v/v) trifluoroacetic acid, 5 μL of sample was injected onto an ACE 5 AQ C-18 column (4.8 mm x 150 mm, 5 μm particle size) with a flow rate of 1 mL/min and the following gradient (Method B): 0 min 30% B, 5 min 100% B, 6 min 100% B, 7 min 30% B, 12 min 30% B. An analytical standard of arcyriaflavin A (Santa-Cruz) was used for comparison.

## Acknowledgements

We would like to thank Professor Mervyn Bibb (John Innes Centre) for personal communication on the *S. venezuelae* arcyriaflavin BGC, which is referenced within the manuscript. HEL was funded by an Imperial College President’s PhD Scholarship. PSF acknowledges funding support from EPSRC (EP/K038648/1, EP/L011573/1). AL was funded as a graduate teaching assistant (University of Kent). LT and SJM acknowledge support from the Leverhulme Trust (RPG-2021-018).

## Data Availability Statement

The data that support the findings of this study are available from the corresponding authors upon reasonable request.

