## Supporting Information for "Biosynthesis of Arcyriaflavin F from *Streptomyces venezuelae* ATCC 10712"

**Table S1.** Oligonucleotides used for cloning of *acfXODCP* or *acfODCP*, and *acfX* expression plasmids.

| **Plasmid** | **Oligonucleotide name** | **Oligonucleotide sequence (5'-3')** |
|---|---|---|
| pAV-$P_{gadph}$-*acfXODCP* | acf_F1_AvrII_F | TGCGAGTATCTGAAAGGGGATACGCCTAGGATGCCGGGAACGTCGACGAC |
| | acf _F1_Rev | GGCGAACAGACCGTGCACGTCG |
| | acf _F2_Fwr | GCCAAGGACCGGATCGACGTGC |
| | acf _F2_Rev | AGAAGTCCAGGGCGTCGGCGTG |
| | acf _F3_Fwr | CTGCTACGACCACGCCGACG |
| | acf _F3_Rev | GCCCGGTTCGGTCACGAAGTGG |
| | acf _F4_Fwr | CCTGGTCCACTTCGTGACCGAACC |
| | acf _F4_PacI_Rev | CTAGAGACTCCTTGCGGATGAGGTTGACGGTTAATTAATCAGGGACGCTCCCGCGAG |
| pAV-$P_{gadph}$-*acfODCP* | acf_X-KO_Fwr | TGCGAGTATCTGAAAGGGGATACGCCTAGGATGCAGACCACCACCGTCATGC |
| pSF1C-A-SP44 | acfX_NdeI_F | ATACATATGCCGGGAACGTCGACGACG |
| | acfX-His6_PacI_R | CACATTAATTAATCAGTGGTGGTGGTGGTGGTGCGCACGGTGTGGGCGGATC |

**Table S2.** NMR characterisation data for Arcyriaflav2in F. Positions are numbered according to Figure 3B.

| Position | $\delta_C$[a] | $\delta_H$[b] (J/Hz) | ROESY | HSQC/ HMQC | HMBC |
|---|---|---|---|---|---|
| **1** | | 10.86 (s) | | | C2, C3, C14, C15 |
| **2** | 171.43 | | | | |
| **3** | 119.43 | | | | |
| **4** | 115.06 | | | | |
| **5** | 129.66 | | | | |
| **6** | | 11.39 (s) | | | C4, C5, C7, C8 |
| **7** | 134.27 | | | | |
| **8** | 122.48 | | | | |
| **9** | 108.82 | 8.40 (d, 2.50) | H25 | C9 | C7, C10, C11 |
| **10** | 151.48 | | | | |
| **11** | 116.32 | 7.02 (dd, 2.51, 8.4) | H12, H25 | C11 | C7, C9, C10 |
| **12** | 112.33 | 7.59 (d, 8.62) | H6, H11 | C12 | C8, C10 |
| **14** | 171.43 | | | | |
| **15** | 119.43 | | | | |
| **16** | 115.06 | | | | |
| **17** | 129.66 | | | | |
| **18** | | 11.39 (s) | | | C16, C17, C19, C20 |
| **19** | 134.27 | | | | |
| **20** | 122.48 | | | | |
| **21** | 108.82 | 8.40 (d, 2.50) | H26 | C21 | C19, C22, C23 |
| **22** | 151.48 | | | | |
| **23** | 116.32 | 7.02 (dd, 2.51, 8.4) | H24, H26 | C23 | C19, C21, C22 |
| **24** | 112.33 | 7.59 (d, 8.62) | H18, H23 | C24 | C20, C22 |
| **25** | | 9.21 (s) | H9, H11 | | |
| **26** | | 9.21 (s) | H21, H23 | | |

[a] Measured at 100 MHz, ppm; [b] Measured at 400 MHz, ppm.

**Figure S1**. Absorbance at 320 nm and EIC of extracts from *S. coelicolor* M1152 recombinant strains grown in MS agar. To detect trace indolocarbazole and arcyriaflavin derivatives, the column was overloaded in comparison to **Figure 2**.

pAV-gapdh – Negative control

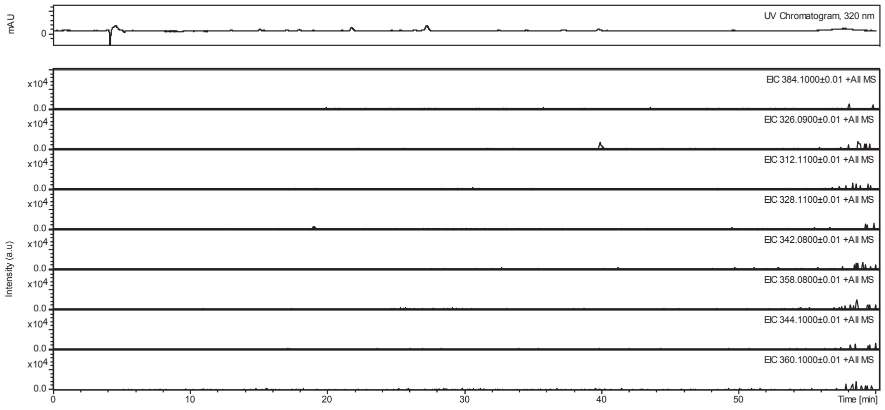


pAV-gapdh-*acfODCP*

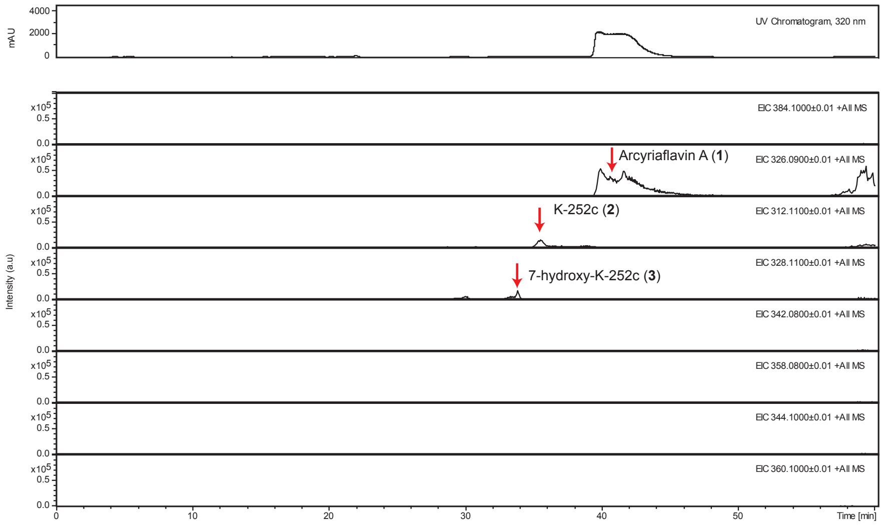


pAV-gadph-*acfXODCP*

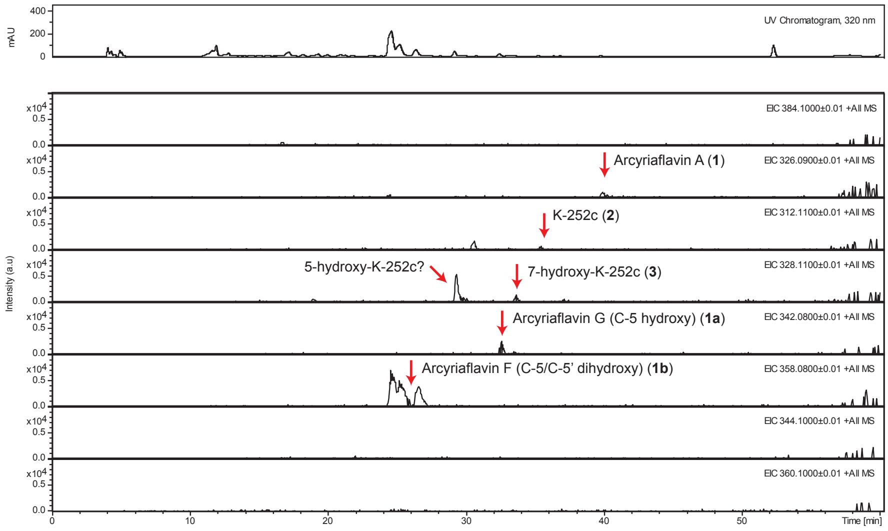

**Figure S2.** NMR spectra of arcyriaflavin F

#### 1H NMR, DMSO-d*6*, 400 MHz

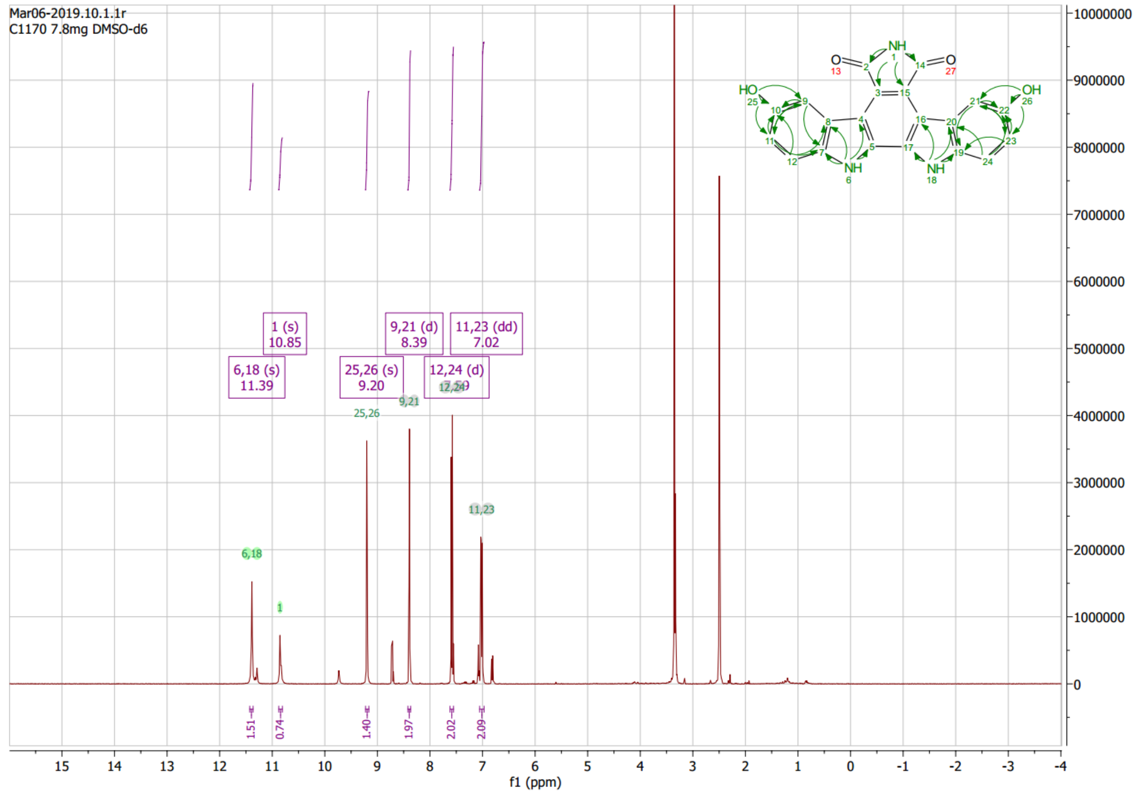


#### 13C NMR, DMSO-d*6*, 100 MHz

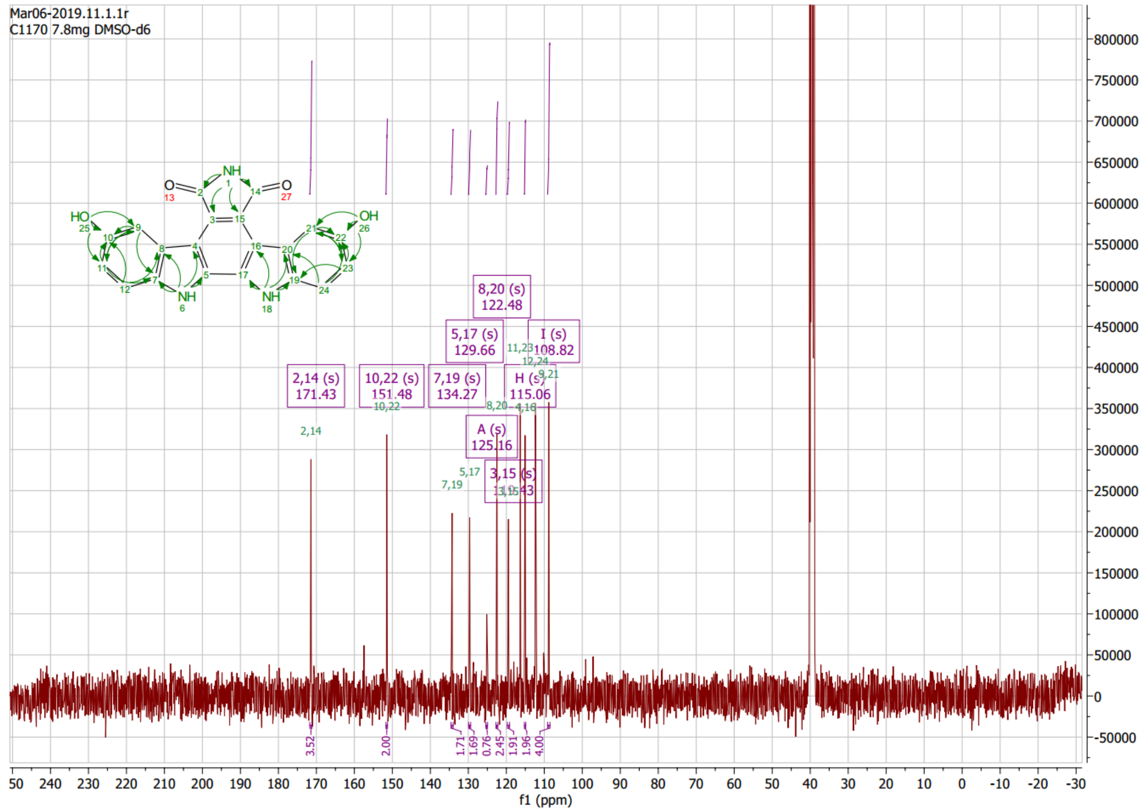

### HMBC

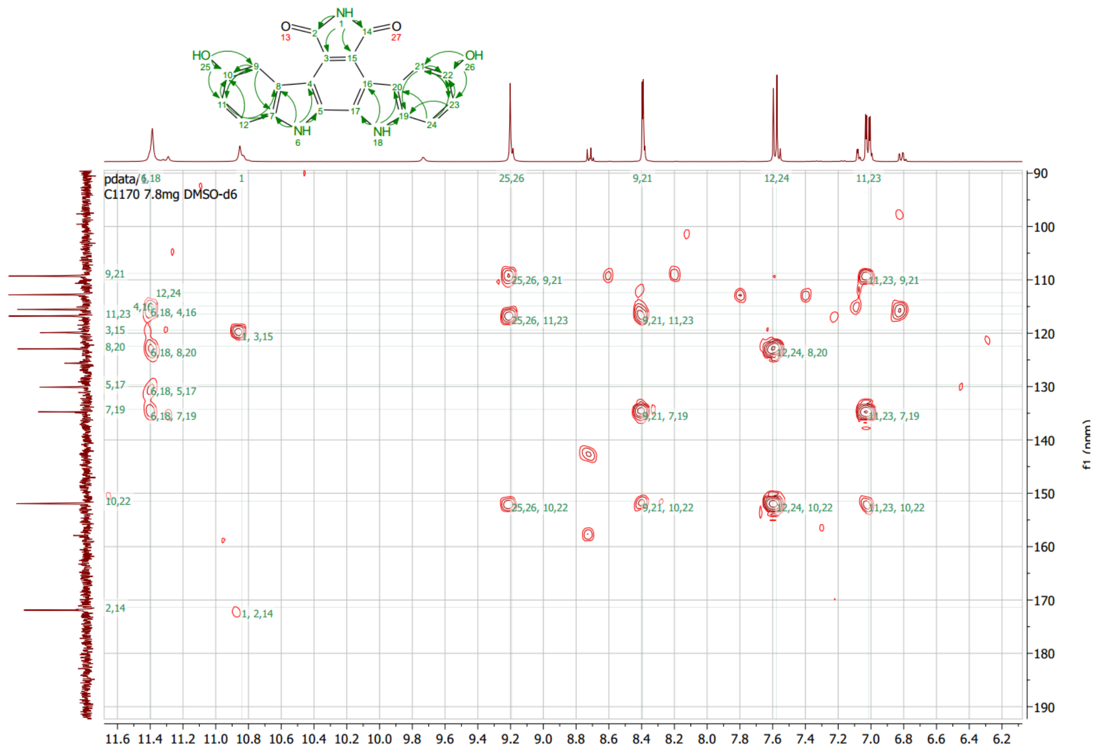
pdata/1
C1170 7.8mg DMSO-d6
f1 (ppm)

### HMQC

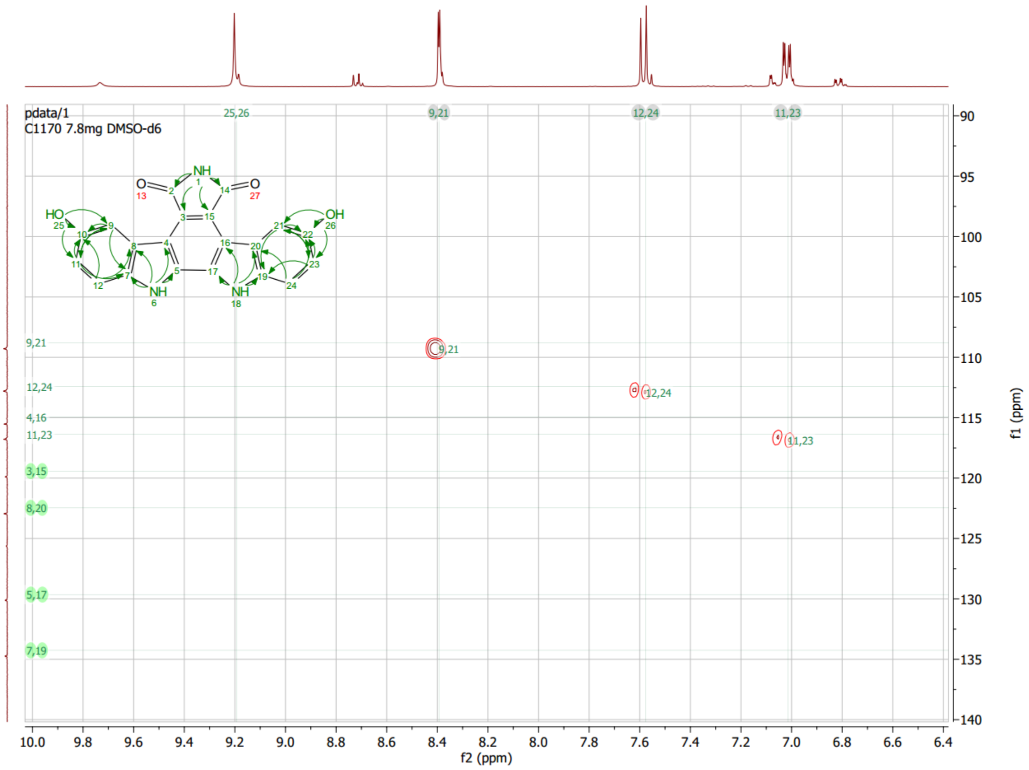
pdata/1
C1170 7.8mg DMSO-d6
f2 (ppm)
f1 (ppm)

#### HSQC

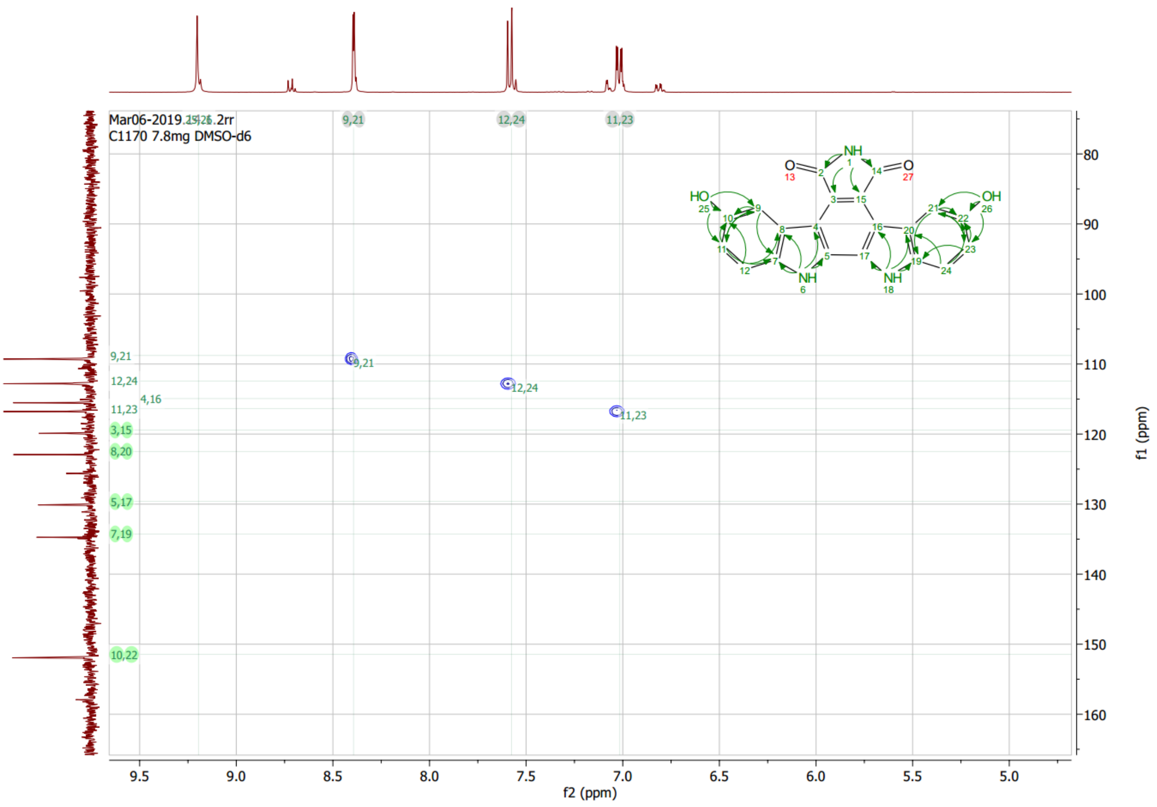
C1170 7.8mg DMSO-d6
9,21
12,24
11,23
4,16
3,15
8,20
5,17
7,19
10,22
f2 (ppm)
f1 (ppm)

#### ROESY

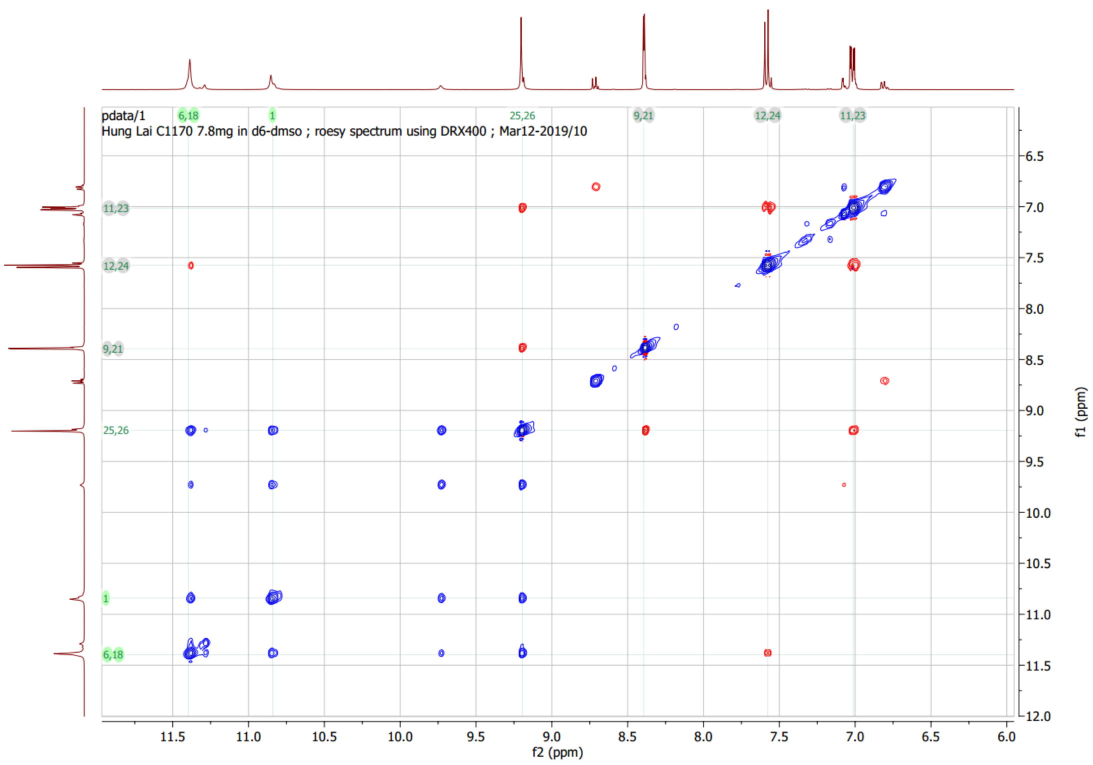
pdata/1
Hung Lai C1170 7.8mg in d6-dmso ; roesy spectrum using DRX400 ; Mar12-2019/10
6,18
1
25,26
9,21
12,24
11,23
f2 (ppm)
f1 (ppm)

#### Superimposed HMBC, HSQC, HMQC

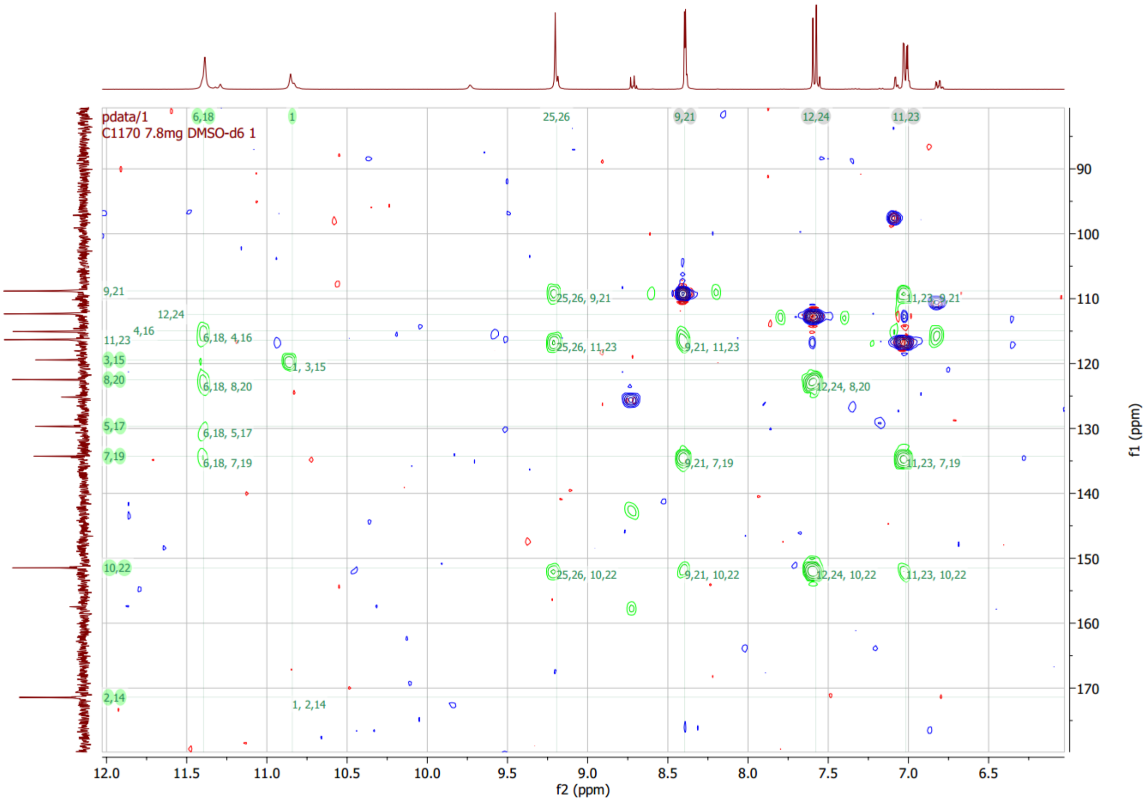

**Figure S3**. AcfX reaction controls and time-course reaction obtained using HPLC Method B

Negative controls. Arcyriaflavin A and F standards (Std) elute at 23.5 min and ~20.7 min.

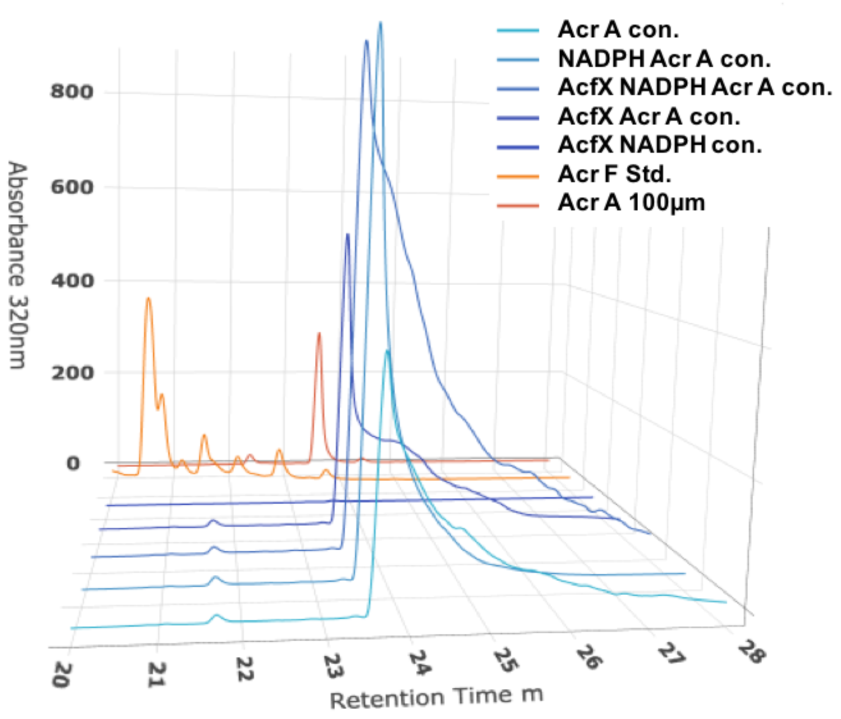


Time course data. Arcyriaflavin A and F elute at 23.5 min and ~20.7 min, respectively on HPLC Method B.

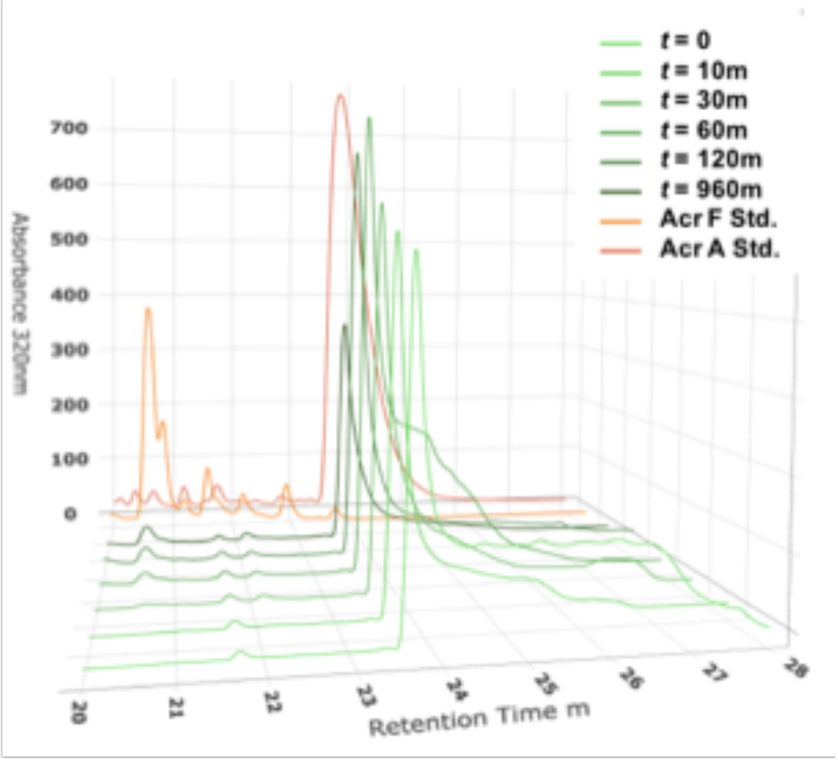
